# PPARgene 2.0: leveraging large language models and multi-omics data for enhanced identification and prediction of PPAR target genes

**DOI:** 10.64898/2025.12.01.691485

**Authors:** Jingxi Qin, Weili Xie, Heng Li, Hao Li, Yucheng Zeng, Yu Luo, Yankai Li, Tianming Sha, Nanping Wang, Yanhui Li, Li Fang

## Abstract

Peroxisome proliferator-activated receptors (PPARs) are ligand-activated transcription factors of the nuclear receptor superfamily. Upon ligand binding, PPARs activate target gene transcription and regulate a variety of important physiological processes such as lipid metabolism, inflammation, wound healing and immune responses. PPARgene is a database that integrates literature-curated and computationally predicted PPAR target genes. It provides gene-level annotations including tissue specificity, species, and supporting PubMed IDs. Computational predictions are generated using a machine learning method that combines PPRE motif analysis with microarray expression data. Here, we introduce PPARgene 2.0, a 10-year update to the original PPARgene database. This update adds 35 newly reported PPARα target genes, 20 PPARβ/δ target genes, and 72 PPARγ target genes, bringing the total number of curated target genes in the database to 337. To retrieve newly reported PPAR target genes from the literature, we used two language models to screen publications after 2016. Candidate papers were then manually reviewed, and verified target genes were added to the updated database. This update also improved the predictive method by expanding the volume of high-throughput gene expression data and incorporating PPAR-related ChIP-seq datasets alongside in silico PPRE analysis. Fivefold cross-validation demonstrated that the new predictive method outperforms the original one. The updated prediction tool is available as part of the PPARgene 2.0 platform. The database is openly accessible at https://www.ppargene.org.

## 1. Introduction

Peroxisome proliferator-activated receptors (PPARs) are ligand-activated transcription factors of the nuclear receptor superfamily. PPARs regulate target gene expression by forming heterodimers with the retinoid X receptor (RXR) and binding to specific DNA sequences known as peroxisome proliferator response elements (PPREs) located in the promoter regions of target genes. Upon ligand binding, conformational changes in PPARs facilitate the recruitment of coactivators and the release of corepressors, thereby modulating transcriptional activity^1^. PPARs play critical roles in the regulation of lipid and glucose metabolism, inflammation, wound healing, and many other pathophysiological processes2–5. Synthetic PPAR ligands, such as fibrates and thiazolidinediones, have been used for clinical treatment of dyslipidemia and type 2 diabetes, respectively^6^.

Identification of PPAR target genes is essential for understanding the molecular mechanisms underlying their diverse biological functions. Numerous PPAR-target genes have been identified since the 1990s^7^. However, the rapid growing of PPAR-related publications each year makes it challenging for researchers to systematically track newly discovered target genes. A comprehensive, well-curated, and freely accessible database of PPAR target genes was essential to support research in this field.

In 2015, we established PPARgene (v1.0)8, the first dedicated database of PPAR target genes. PPARgene (v1.0) offers a manually curated collection of experimentally validated targets, complete with detailed annotations including PPAR subtypes, tissue specificity, species, and supporting literature. In addition, it includes predicted PPAR target genes based on high-throughput gene expression data and PPRE motif matching.

Over the past decade, PPAR-related research has expanded dramatically, with over 20,000 new studies published, reflecting the growing interest in PPAR biology. In parallel, advances in high-throughput sequencing technologies have generated vast amounts of transcriptomic and epigenetic data—including RNA-seq, ChIP-seq, and ATAC-seq—providing unprecedented opportunities to explore PPAR-mediated gene regulation at a systems level. These developments underscore the urgent need to update PPARgene (v1.0) to incorporate the latest experimental findings and leverage newly available omics data.

In this study, we present PPARgene version 2.0, a major update of the database after a decade, featuring substantially expanded content and an enhanced predictive framework. To identify novel PPAR target genes, we employed two large language models (LLMs) to systematically screen publications released after 2015. All candidate genes identified by the LLMs underwent rigorous manual curation prior to database inclusion.

To improve the accuracy of target gene prediction, we constructed a multi-omics dataset that integrates transcriptional evidence (microarray and RNA-seq), experimental PPAR binding information (ChIP-seq), and in silico PPRE motif predictions. The updated machine learning model, trained on this comprehensive data, demonstrates significantly improved predictive performance compared to the original v1.0 model.

Overall, PPARgene 2.0 provides a more comprehensive and reliable resource for studying PPARs and their downstream regulatory networks. The database is freely accessible at www.ppargene.org.

## 2. Methods

### 2.1. Data Collection

#### 2.1.1. Collection of Experimentally Verified PPAR Target Genes

To incorporate newly discovered PPAR target genes identified after 2016 into the PPARgene database, we examined abstracts of all articles indexed in PubMed between January 1, 2016, and June 20, 2024. Two language models, Bioformer9 and GPT-410, were employed to screen these articles independently. Bioformer is a BERT-like language model for biomedical text mining. For both language models, abstracts were initially filtered using regular expressions to retain only those containing PPAR-related terms. The resulting subset of abstracts was used as input for model evaluation.

We employed a BERT-based text classification approach utilizing the Bioformer model for screening. To facilitate this, a balanced fine-tuning dataset was constructed comprising abstracts from the first version of the PPARgene database as positive samples and an equivalent number of randomly selected pre-2016 PubMed abstracts as negative samples. The Bioformer model was fine-tuned on this dataset and subsequently applied to classify the collected abstracts, retaining only those predicted as positive.

For the GPT-based screening, we employed a two-step procedure. First, the GPT-4o-mini-2024-07-18 model was used to perform an initial assessment of whether an abstract was related to PPAR target genes. Abstracts for which the model output was “Yes” were retained. In the second step, these abstracts were evaluated by the more advanced GPT-4o-2024-08-06 model. This model was prompted, using carefully engineered chain-of-thought instructions (see Supplementary Materials for details), to specifically identify articles that experimentally reported PPAR target genes. Abstracts receiving a final response of “Yes” were kept for further analysis.

Finally, the results from both Bioformer and GPT screenings were manually reviewed to extract newly reported PPAR target genes.

#### 2.1.2. Collection of PPAR-Relevant Microarray & RNA-seq Data Sets

PPAR-relevant microarray and RNA-seq data sets were acquired by searching the GEO database^11^ using the key words “PPAR”, “PPAR alpha”, “PPAR beta”, “PPAR delta”, “PPAR gamma”, or “Peroxisome Proliferator-Activated Receptor”. In total, we curated 68 microarray and 15 RNA-seq data sets in which PPAR isoforms were activated, overexpressed, suppressed, knockout or knockdown.

#### 2.1.3. Collection of PPAR ChIP-seq and ATAC-seq Data Sets

To investigate the genomic binding sites of Peroxisome Proliferator-Activated Receptors (PPARs), we obtained IDR-thresholded ChIP-seq peak call sets for PPARα, PPARδ, and PPARγ from the ENCODE portal (accession numbers: ENCFF708MEQ, ENCFF893YKY, and ENCFF567JIZ, respectively)^12–14^. Corresponding IDR-thresholded ATAC-seq peak sets from the same cell lines were also retrieved to assess chromatin accessibility (accession numbers: ENCFF855CQR, ENCFF948AFM, and ENCFF935GLR, respectively). All ENCODE datasets were based on the GRCh38 reference genome (ENCODE4, v1.6.1).

Additionally, we utilized a second set of PPAR ChIP-seq data from the ChIP-Atlas database^15^. Using its Target Genes Tool, we identified genes with a PPAR binding peak, called by MACS2, located within a ±5 kb window of their transcription start site (TSS).

### 2.2. Feature Extraction

#### 2.2.1. Transcriptomic Evidence (TE)

To assemble high-throughput gene expression evidence for Peroxisome Proliferator-Activated Receptor (PPAR) target gene interactions, we curated a collection of microarray and RNA-seq datasets. These datasets encompassed various experimental perturbations of PPARα, PPARγ, and PPARδ, including their activation, overexpression, knockout, or knockdown.

The data comprised 68 microarray experiments from Homo sapiens, Mus musculus, Macaca fascicularis, and Rattus norvegicus. Raw microarray data were processed using the Bioconductor package16. To harmonize gene identifiers across species, all genes were mapped to their human orthologs based on the Affymetrix Human Genome U133 Plus 2.0 Array (GPL570) annotation. Only genes present on this platform were retained for subsequent analysis.

Additionally, 15 RNA-seq datasets from Homo sapiens and Mus musculus were included. We obtained normalized expression matrices (e.g., TPM or raw counts) from the Gene Expression Omnibus (GEO) and performed differential gene expression analysis using DESeq217. For these datasets, gene annotation was standardized against the human GPL11154 platform, again retaining only the genes present in this reference.

In total, 83 differential gene expression (DGE) experiments were integrated. A final, comprehensive gene set was created by taking the intersection of genes present in both the harmonized microarray (GPL570) and RNA-seq (GPL11154) datasets. To ensure data completeness, missing log-fold change (logFC) values within any given experiment were imputed using the median logFC of that experiment.

To summarize each gene’s differential expression signature across the 83 experiments, we engineered two sets of proportion-based features from their log fold change (logFC) values.

1. Threshold-Based Features. We calculated the proportion of experiments in which a gene’s logFC exceeded specific positive or negative thresholds. This resulted in six features per gene, corresponding to the proportions for logFC > 1.5, logFC < -1.5, logFC > 1.0, logFC < -1.0, logFC > 0.5, and logFC < -0.5. For instance, a gene exhibiting logFC > 1.5 in 23 of the 83 experiments received a feature value of 23/83 (≈ 0.277).
2. Rank-Based Feature. We computed the proportion of experiments where a gene ranked among the top 2% of all genes based on its logFC. This required ranking all genes by logFC within each experiment individually beforehand.

#### 2.2.2. In silico PPRE Evidence (ISPE)

To characterize the presence of conserved PPAR response elements (PPREs), we introduced a binary feature indicating whether a gene harbors any predicted PPREs within the ±5 kb region flanking the transcription start site (TSS) in at least one of the three species: human, mouse, or rat. A value of 1 was assigned if PPREs were identified in this region in any species; otherwise, the value was 0.

The prediction of PPREs was based on previously published sequence-level motif scanning approaches. PPREs were predicted using position weight matrix (PWM) models derived from the PPARγ-RXRα heterodimer binding motif, without distinguishing among PPAR subtypes due to their shared core consensus sequence. Motif scanning, PWM score computation, and background correction followed standard methods as described in our previous study^8^.

#### 2.2.3. PPAR Binding Evidence (PBE)

To construct features representing direct transcription factor (TF) binding, we processed the ChIP-seq and ATAC-seq datasets described in Section 2.1.3.

For the ENCODE data, we defined a feature based on peak signal strength. For each gene, we first identified PPAR ChIP-seq peaks within the ±5 kb region flanking its transcription start site (TSS) that overlapped with ATAC-seq signals from its corresponding cell line. From this set of co-localized peaks, we retained only those with a q-value less than 0.1. If multiple significant peaks remained for a gene, we selected the single peak with the highest signalValue, which represents the average enrichment strength. This final signalValue was then log-transformed and standardized for use as a feature in our downstream analysis.

For the ChIP-Atlas data, we created a feature based on binding frequency. Using the Target Genes tool, we retrieved genes with ChIP-seq peaks for human PPARA, PPARD, and PPARG (hg38) and mouse Ppara, Ppard, and Pparg (mm10) within a ±5 kb window of their TSS. A gene was considered bound in an experiment if it had a peak with a MACS2 score of 250 or greater. For each PPAR subtype, the feature was calculated as the proportion of experiments showing binding, i.e., the number of experiments meeting the score threshold divided by the total number of available experiments for that TF.

### 2.3. Model Training and Evaluation

#### 2.3.1. Training Sets for the Prediction Model

We used experimentally validated target genes from the PPARgene 2.0 database as the positive training set. Given that PPAR exists as three distinct isoforms (PPARα, PPARβ/δ, and PPARγ), we constructed four separate classification models: a general pan-PPAR model and three isoform-specific models. For the isoform-specific models, we engineered features using corresponding ChIP-seq data from the ENCODE and ChIP-Atlas databases.

Constructing the negative training sets for the isoform-specific models presented a notable challenge, as it is difficult to definitively confirm that a gene is not a PPAR target under all possible biological conditions or in any cell type. Furthermore, due to the high similarity in DNA-binding specificity among PPAR isoforms, the absence of reported evidence for one isoform does not preclude a gene from being a target of another. We therefore adopted a conservative strategy for building the isoform-specific negative sets by excluding all experimentally verified PPAR target genes, irrespective of their isoform. The negative samples were generated by randomly selecting an equal number of genes from a background gene pool, from which pseudogenes and genes with LOC-prefixed symbols had also been removed.

To mitigate potential sampling bias, we repeated this negative sampling process 100 times. Each model was subsequently trained 100 times, pairing the static positive set with each of the 100 independently generated negative sets.

#### 2.3.2. Logistic Regression Classifier

We trained four binary logistic regression models to predict PPAR target genes: one general pan-PPAR model and three isoform-specific models (PPARα, PPARβ/δ, and PPARγ), using the features described above. Let *pi* be the probability that the *i-*th gene is a PPAR target gene. The logistic regression model is

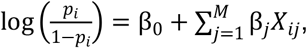

Where *βj* is the regression coefficient of the feature *Xij*. The logistic regression model was implemented using the LogisticRegression class from the scikit-learn Python library^18^.

#### 2.3.3. Performance Evaluation

We used 5-fold cross validation to evaluate the performance of the logistic regression model. In each round, 20% of the samples were left out as the test data and the remaining were the training data. To convert predicted probabilities into binary labels, we applied a classification threshold of 0.4. Precision, recall, and F1 score were used to evaluate the performance of the classifier. Precision, recall, and F1 were calculated as

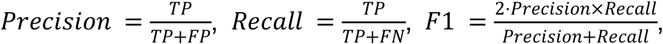

where TP is the number of true positives, FP is the number of false positives, and FN is the number of false negatives. We also calculated AUC, the area under the receiver operating characteristic (ROC) curve, using the roc_auc_score function from scikit-learn. Because negative data sets were obtained by 100 random samplings, the medians of precisions, recalls, F1 scores, and AUCs of the 100 training results were used.

### 2.4. Genome-Wide Prediction of PPAR Target Genes

We trained four logistic regression models (a general pan-PPAR model and three isoform-specific models) using a 1:5 ratio of positive to negative samples and the previously described feature set. The classification thresholds for predicting PPAR target genes were determined by selecting the probability values corresponding to precisions of 0.8, 0.65, and 0.5. Genes with predicted probabilities above the threshold corresponding to precision 0.8 were classified as high-confidence targets; those with probabilities between the thresholds for precisions 0.65 and 0.8 as medium-confidence targets; and those with probabilities between the thresholds for precisions 0.5 and 0.65 as low-confidence targets. Genes with predicted probabilities below the threshold corresponding to precision 0.5 were predicted as non-targets.

### 2.5. Web Server

PPARgene 2.0 is implemented using Flask with a Bootstrap frontend and an SQLite backend. It is publicly available at https://www.ppargene.org/.

## 3. Results and Discussion

### 3.1. Experimentally Verified PPAR Target Genes

The update to the PPARgene database incorporates an additional 112 experimentally verified PPAR target genes, bringing the total to 337. This expansion is the result of a comprehensive literature search and screening pipeline applied to articles published between January 1, 2016, and June 20, 2024.

Our screening process began with an initial pool of 23,746 PPAR-related abstracts. The language model-based workflow (as described in Section 2.1.1) successfully narrowed this collection down to 441 high-confidence articles that were prioritized for manual curation. Following strict manual validation of the experimental evidence within these articles, the number of verified targets in the PPARgene 2.0 database increased from 83 to 118 for PPARα, from 83 to 103 for PPARβ/δ, and from 104 to 176 for PPARγ.

### 3.2. Development of Logistic Regression Models to Predict PPAR Target Genes

To predict novel PPAR target genes, we trained logistic regression models using experimentally validated targets as positive samples and an equal number of randomly sampled background genes as negative controls. Given that the three PPAR subtypes recognize a conserved core sequence and frequently share targets, all validated genes were pooled for training without subtype distinction.

We initially trained the model using a single feature—transcriptomic evidence (TE)—as fold-change data has proven effective in prior gene prediction studies8. Five-fold cross-validation (Table 1) yielded median precision, recall, F1 score, and AUC values of 0.710, 0.800, 0.754, and 0.820, respectively. Notably, compared to PPARgene v1.0—which incorporates both transcriptomic and PPAR binding evidence—our model matched its AUC while delivering higher recall and F1 scores. These results suggest that the inclusion of a larger number of high-throughput datasets in PPARgene v2 significantly improves predictive performance.

**Table 1.** Performances of logistic regression models trained on different features.

| PPAR Subtype | Features | Precision | Recall | F1 | AUC |
| --- | --- | --- | --- | --- | --- |
| PPARs (v2.0) | TE | 0.710 | 0.800 | 0.754 | 0.820 |
|  | PBE | 0.646 | 0.817 | 0.717 | 0.766 |
|  | ISPE | 0.570 | 0.915 | 0.703 | 0.610 |
|  | TE + PBE + ISPE | 0.718 | 0.817 | 0.763 | 0.843 |
| PPARs (v1.0) | CPS + HTE | 0.83 | 0.59 | 0.69 | 0.82 |
| PPARG (v2.0) | TE | 0.698 | 0.733 | 0.714 | 0.787 |
|  | PBE | 0.509 | 0.967 | 0.667 | 0.758 |
|  | ISPE | 0.569 | 0.900 | 0.701 | 0.604 |
|  | TE + PBE + ISPE | 0.696 | 0.733 | 0.721 | 0.811 |
| PPARA (v2.0) | TE | 0.783 | 0.810 | 0.796 | 0.876 |
|  | PBE | 0.606 | 0.952 | 0.741 | 0.837 |
|  | ISPE | 0.576 | 0.950 | 0.717 | 0.619 |
|  | TE + PBE + ISPE | 0.792 | 0.810 | 0.800 | 0.891 |
| PPARD (v2.0) | TE | 0.688 | 0.550 | 0.606 | 0.680 |
|  | PBE | 0.500 | 1.000 | 0.667 | 0.536 |
|  | ISPE | 0.571 | 0.950 | 0.706 | 0.625 |
|  | TE + PBE + ISPE | 0.576 | 0.900 | 0.691 | 0.730 |

TE: Transcriptomic Evidence; PBE: PPAR Binding Evidence; ISPE: *In silico* PPRE Evidence; CPS: conserved PPRE score; HTE: high throughput evidence.

We next trained a model based solely on in silico PPRE evidence (ISPE), a binary feature indicating the presence of predicted PPAR response elements (PPREs) conserved across mouse, rat, and human genomes within ±5 kb of the transcription start site. This model attained a precision of 0.570, recall of 0.915, F1 score of 0.703, and AUC of 0.610 (Table 1).

Since ChIP-seq provides direct experimental evidence of PPAR binding events—offering a key advantage over purely in silico predictions—we constructed a PPAR Binding Evidence (PBE) feature using publicly available ChIP-seq data from ENCODE and ChIP-Atlas (as described in Section 2.2.3). The inclusion of PBE features led to improved model performance, achieving a precision of 0.646, recall of 0.817, F1 score of 0.717, and an AUC of 0.766 (Table 1). This represents a clear enhancement over the model relying solely on in silico PPRE evidence.

Finally, we built an integrated logistic regression model combining transcriptomic evidence (TE), PPAR binding evidence (PBE), and in silico PPRE evidence (ISPE). This combined model exhibited median precision, recall, F1 score, and AUC values of 0.718, 0.817, 0.763, and 0.843, respectively, outperforming the best-performing model from PPARgene v1.0 (Table 1 and Figure 1). In addition, we performed an ablation analysis to evaluate the contribution of each feature group. All tested features contributed to AUROC to varying degrees, with transcriptomic evidence (TE) having the most substantial impact. Detailed results of the ablation analysis are provided in Figure 1 and Supplementary Table 1.

**Figure 1.**
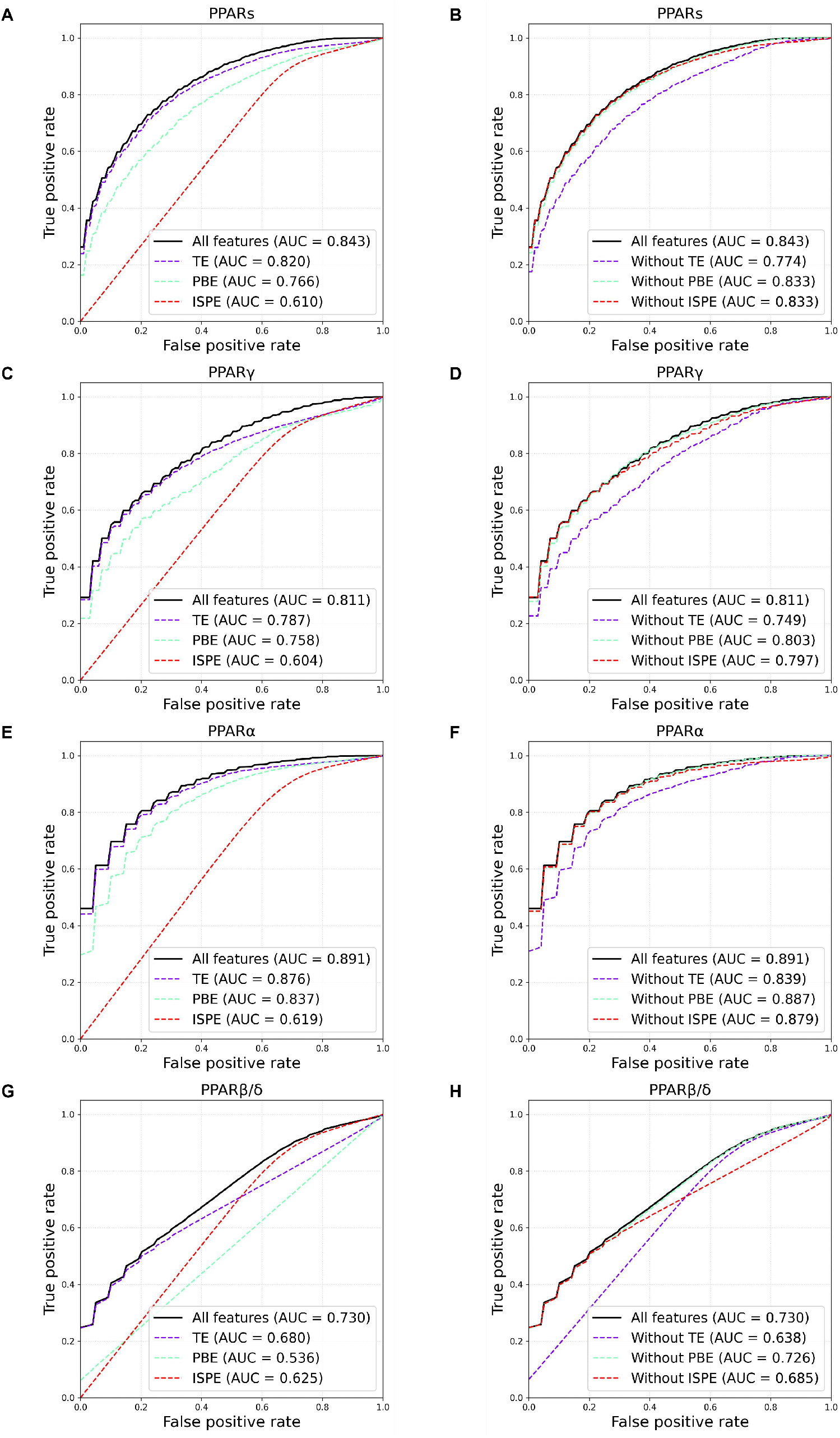
ROC curves for logistic regression models trained on different features. (A, C, E, G) Model performance using all features or individual features. ROC curves compare the predictive accuracy of pan-subtype and subtype-specific models when trained exclusively on a single feature type. (B, D, F, H) Overall and subtype-specific model performance with ablation analysis. ROC curves depict the performance of pan-subtype (PPARs) and subtype-specific (PPARγ, PPARα, PPARβ/δ) models in predicting target genes. Each model’s curve is presented alongside results from its corresponding ablation analysis. TE: Transcriptomic Evidence; ISPE: *In silico* PPRE Evidence; PBE: PPAR Binding Evidence

### 3.3. Isoform-specific Prediction of PPAR Target Genes

We developed separate models for each of the three PPAR isoforms using isoform-specific features (i.e., TE and PBE). Given the high similarity of their core motifs, the same algorithm was employed for in silico prediction across all three isoforms. The PPARα model exhibited the best performance, which may be explained by the higher quality and consistency of its training data, with median precision, recall, F1-score, and AUC values of 0.792, 0.810, 0.800, and 0.891, respectively. In contrast, the PPARδ model showed comparatively lower performance, likely due to the limited amount of available data; its corresponding median precision, recall, F1-score, and AUC were 0.576, 0.900, 0.691, and 0.730 (Table 1). The results of the ablation study are shown in Supplementary Table 1.

### 3.4. Genome-Wide Prediction of PPAR Target Genes

The general model predicted 2,533 target genes (420 high-confidence, 753 medium-confidence, 1,360 low-confidence). The isoform-specific analysis revealed 3,432 targets for PPARα, 2,385 for PPARγ, and 1,490 for PPARδ, with varying confidence distributions detailed in Figure 2A. Specifically, high-confidence predictions numbered 582, 309, and 264 for the α, γ, and δ isoforms, respectively. The confidence breakdown and model intersections are further visualized in Figure 2 (pie charts and UpSet plot). All predicted targets from both model types are cataloged on the PPARgene 2.0 website.

**Figure 2.**
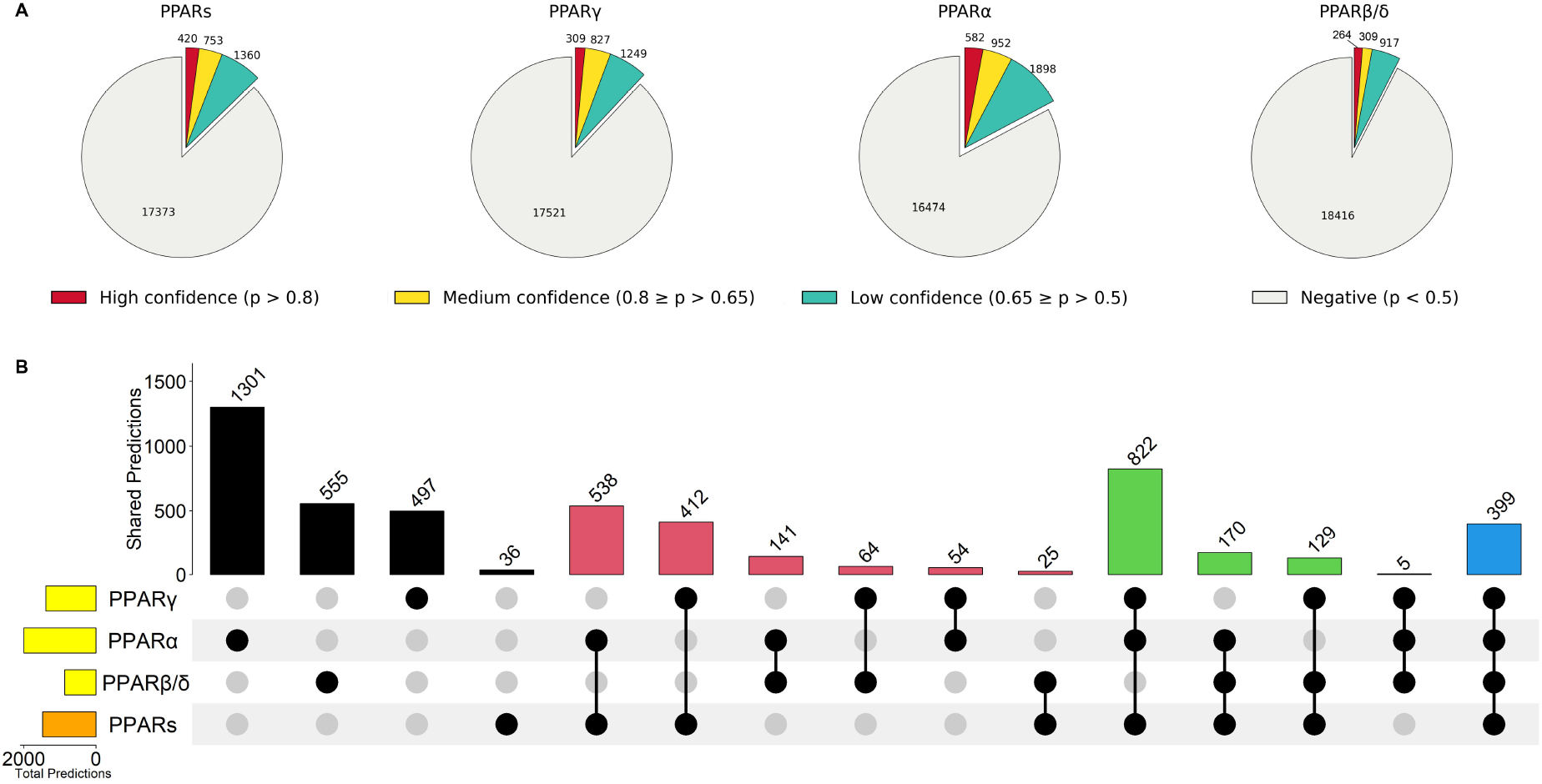
Genome-wide Prediction of PPAR Target Genes. (A)Prediction results and confidence classification of PPAR target genes. This figure presents the predictions from the pan-PPAR and subtype-specific models. Genes are classified as high-, medium-, or low-confidence targets based on probability thresholds set to precision levels of 0.8, 0.65, and 0.5, as derived from logistic regression models trained with a 1:5 positive-to-negative sample ratio. (B)Overlap of predicted PPAR target genes across models. This UpSet plot visualizes the intersections of target genes predicted by the pan-PPAR and subtype-specific models, showing the number of genes uniquely predicted by a single model, as well as those shared by two, three, or all models.

### 4. Querying the Database

The PPARgene 2.0 database is composed of two modules: one is for querying experimentally verified target genes and the other is for querying computationally predicted target genes.

#### 4.1. Experimentally Verified Target Genes

We provide users two ways to query the experimentally verified target genes. First, users can browse the results by selecting the PPAR subtype. PPARgene 2.0 will return a table of matched entries. Users can also submit a specific gene symbol. The provided results contain the following items: PPAR subtype, gene symbol, species, tissue/cell types, regulation direction, and reference PubMed IDs.

#### 4.2. Computationally Predicted Target Genes

Users can retrieve the prediction results by querying the gene symbol. If the gene is predicted as a PPAR target gene, the query will return a *p* value (the probability of being a PPAR target gene) with a confidence level (high, medium or low). On the website, this probability is referred to and can be queried as the “PPAR Target Score”. A larger PPAR Target Score means a higher confidence. Transcriptomic evidence, PPAR binding evidence and existence of putative PPREs were listed to support the prediction.

#### 4.3. Downloadable Files

Users can download data sets of experimentally verified PPAR target genes as well as computationally predicted target genes. We also provide hyperlinks for downloading the high throughput experimental data sets curated for our prediction model.

### 5. Future Extensions

In this release of the PPARgene 2.0 database, we have focused on the curation and isoform-specific prediction of PPAR target genes. In future updates, we will incorporate additional epigenomic features (e.g., histone modifications), interspecies conservation data, and chromatin contact information (e.g., Hi-C) to further improve prediction accuracy and biological interpretability. We also plan to expand the database to include additional nuclear receptors beyond PPARs and to develop dedicated prediction models for each nuclear receptor family.

## 6. Conclusion

In this study, we present PPARgene 2.0, a major update to the original PPARgene database released a decade ago. This new version adds 35 newly identified PPARα target genes, 20 PPARβ/δ target genes, and 72 PPARγ target genes, bringing the total number of experimentally validated PPAR targets to 337—a 60% increase over the previous release.

To enable reliable prediction of PPAR target genes, we have integrated an extensive collection of high-throughput datasets, including 85 gene expression profiles (from microarray and RNA-seq experiments) along with binding profiles from ChIP-seq and ATAC-seq. Our prediction algorithm incorporates a comprehensive set of features, such as in silico PPRE motif prediction, high-throughput gene expression data, and PPAR binding evidence from ChIP-seq and ATAC-seq. Furthermore, we developed isoform-specific machine learning models for PPARα, PPARβ/δ, and PPARγ to predict potential PPAR target genes across the human genome. We believe that PPARgene 2.0 will serve as a valuable and up-to-date resource for the PPAR research community.

## Acknowledgments

This work was funded by grants from the National Natural Science Foundation of China (32300546), GuangDong Basic and Applied Basic Research Foundation (2025A1515012992) and Sun Yat-sen University College Student Innovation and Entrepreneurship Training Program (Grant No. 20252763)

## Notes

https://www.ppargene.org

### Competing Interest Statement

The authors have declared no competing interest.

## References

1. Berger, J. & Moller, D. E. The mechanisms of action of PPARs. Annu Rev Med 53, 409–435 (2002).

2. Fan, Y. et al. Suppression of pro-inflammatory adhesion molecules by PPAR-delta in human vascular endothelial cells. Arterioscler Thromb Vasc Biol 28, 315–321 (2008).

3. Harman, F. S. et al. Peroxisome proliferator-activated receptor-delta attenuates colon carcinogenesis. Nat Med 10, 481–483 (2004).

4. Wang, N. et al. Constitutive activation of peroxisome proliferator-activated receptor-gamma suppresses pro-inflammatory adhesion molecules in human vascular endothelial cells. J Biol Chem 277, 34176–34181 (2002).

5. Wang, Y.-X. PPARs: diverse regulators in energy metabolism and metabolic diseases. Cell Res 20, 124–137 (2010).

6. Staels, B. & Fruchart, J.-C. Therapeutic roles of peroxisome proliferator-activated receptor agonists. Diabetes 54, 2460–2470 (2005).

7. Rakhshandehroo, M., Knoch, B., Müller, M. & Kersten, S. Peroxisome proliferator-activated receptor alpha target genes. PPAR Res 2010, 612089 (2010).

8. Fang, L. et al. PPARgene: A Database of Experimentally Verified and Computationally Predicted PPAR Target Genes. PPAR Res 2016, 6042162 (2016).

9. Fang, L., Chen, Q., Wei, C.-H., Lu, Z. & Wang, K. Bioformer: an efficient transformer language model for biomedical text mining. ArXiv 2302.01588v1 (2023).

10. OpenAI et al. GPT-4 Technical Report. Preprint at 10.48550/arXiv.2303.08774 (2024).

11. Clough, E. et al. NCBI GEO: archive for gene expression and epigenomics data sets: 23-year update. Nucleic Acids Res 52, D138–D144 (2024).

12. ENCODE Project Consortium. An integrated encyclopedia of DNA elements in the human genome. Nature 489, 57–74 (2012).

13. Luo, Y. et al. New developments on the Encyclopedia of DNA Elements (ENCODE) data portal. Nucleic Acids Res 48, D882–D889 (2020).

14. Bc, H. et al. The ENCODE Uniform Analysis Pipelines. bioRxiv : the preprint server for biology https://doi.org/10.1101/2023.04.04.535623 (2023) doi:10.1101/2023.04.04.535623.

15. Zou, Z., Ohta, T. & Oki, S. ChIP-Atlas 3.0: a data-mining suite to explore chromosome architecture together with large-scale regulome data. Nucleic Acids Res 52, W45–W53 (2024).

16. W, H. et al. Orchestrating high-throughput genomic analysis with Bioconductor. Nature methods 12, (2015).

17. Love, M. I., Huber, W. & Anders, S. Moderated estimation of fold change and dispersion for RNA-seq data with DESeq2. Genome Biol 15, 550 (2014).

18. Pedregosa, F. et al. Scikit-learn: Machine Learning in Python. arXiv.org https://arxiv.org/abs/1201.0490v4 (2012).

