## Supplementary Material for "PPARgene 2.0: leveraging large language models and multi-omics data for enhanced identification and prediction of PPAR target genes"

**ADDITIONAL FILE 1.**

**Prompts used when calling gpt-4o-mini-2024-07-18**

Based on the above text, is there any gene identified as a direct target gene of PPAR? Please simply answer the gene name. Answer "None" if no gene was identified as a target gene of PPAR.

**Prompts used when calling gpt-4o-2024-08-06**

1. Based on the above text, does it describe a target gene of PPAR? Answer "Yes" and target gene's name or "No".

2. Based on the above text, does it describe the process to discover PPAR's target gene? Answer "Yes" or "No".

3. Based on the above text, does it describe the target gene's expression is dependent on PPAR? Answer "Yes" or "No".

4. Based on the above text, does it describe the target gene's expression including level of mRNA or protein is changed because of the activation or deletion of PPAR?

5. Based on the above text, does it mention any experiment that prove any part of the target gene that binds PPAR? Answer "Yes" or "No".

6. Based on the above text and question 1 to 5, is the text describe a target gene of PPAR that has been found by experiment, or just describe a target gene of PPAR without doing any experiment to prove it? Answer "Yes" for the first choice or "No" the other.

Table 1. Performances of logistic regression models in ablation study.

| **PPAR Subtype** | **Features** | **Precision** | **Recall** | **F1** | **AUC** |
| --- | --- | --- | --- | --- | --- |
| PPARs (v2.0) | TE + PBE + ISPE | 0.718 | 0.817 | 0.763 | 0.843 |
|  | w/o TE | 0.585 | 0.898 | 0.712 | 0.774 |
|  | w/o PBE | 0.707 | 0.800 | 0.756 | 0.833 |
|  | w/o ISPE | 0.721 | 0.814 | 0.760 | 0.833 |
| PPARG (v2.0) | TE + PBE + ISPE | 0.696 | 0.733 | 0.721 | 0.811 |
|  | w/o TE | 0.564 | 0.900 | 0.699 | 0.749 |
|  | w/o PBE | 0.700 | 0.767 | 0.730 | 0.803 |
|  | w/o ISPE | 0.700 | 0.733 | 0.717 | 0.797 |
| PPARA (v2.0) | TE + PBE + ISPE | 0.792 | 0.810 | 0.800 | 0.891 |
|  | w/o TE | 0.576 | 0.950 | 0.717 | 0.839 |
|  | w/o PBE | 0.789 | 0.810 | 0.800 | 0.887 |
|  | w/o ISPE | 0.789 | 0.810 | 0.800 | 0.879 |
| PPARD (v2.0) | TE + PBE + ISPE | 0.576 | 0.900 | 0.691 | 0.730 |
|  | w/o TE | 0.571 | 0.950 | 0.706 | 0.638 |
|  | w/o PBE | 0.576 | 0.900 | 0.691 | 0.726 |
|  | w/o ISPE | 0.688 | 0.550 | 0.606 | 0.685 |

TE: Transcriptomic Evidence; PBE: PPAR Binding Evidence; ISPE: *In silico* PPRE Evidence
